# Transcriptomic Response to Nitrogen Availability Highlights Signatures of Adaptive Plasticity During Tetraploid Wheat Domestication

**DOI:** 10.1101/2023.08.31.555682

**Authors:** Alice Pieri, Romina Beleggia, Tania Gioia, Hao Tong, Valerio di Vittori, Giulia Frascarelli, Elena Bitocchi, Laura Nanni, Elisa Bellucci, Fabio Fiorani, Nicola Pecchioni, Concetta De Quattro, Antonina Rita Limongi, Pasquale De Vita, Marzia Rossato, Ulrich Schurr, Jacques L. David, Zoran Nikoloski, Roberto Papa

**Author notes:** Corresponding authors contact: Roberto Papa; **Romina Beleggia**.

## Abstract

The domestication of crops, with the development of the agroecosystems, is associated with major environmental changes and represent a model to test the role of phenotypic plasticity. Here we investigated 32 genotypes representing key stages of tetraploid wheat domestication. We developed a dedicated pipeline combining RNA-Seq data, estimates of evolvability and *Q_ST_* to characterize the plasticity of gene expression and identify signatures of selection under different nitrogen conditions. The analysis of gene expression diversity showed contrasting results between primary and secondary domestication in relation to nitrogen availability. Indeed, nitrogen triggered the expression of twice the number of genes in durum wheat compared to emmer and wild emmer. *Q_ST_* distributions and *Q_ST_–F_ST_* comparisons revealed distinct selection signatures at each domestication stage. While primary domestication affected the expression of genes involved in biotic interactions, secondary domestication was associated with changes in expression of genes involved in metabolism of amino acids, particularly lysine. Selection signatures were found also in differentially expressed genes, specifically involved in nitrogen metabolism, such as *glutamate dehydrogenase*. Overall, our findings show that nitrogen availability had a pivotal role during the domestication and adaptive responses of a major food crop, with varying effects across different traits and growth conditions.

## Introduction

Domestication influences the genetic diversity of animals and plants as they adapt to agroecosystems, and undergo selection to meet human preferences and needs. This process is typically associated with the genome-wide loss of nucleotide diversity due to the combined consequences of selection and genetic drift, which is known as the domestication bottleneck. The loss of genetic diversity has been documented in many domesticated species by comparing them with wild relatives (Bitocchi et al., 2017). A parallel effect is the reprogramming of gene expression and the loss of expression diversity, which was first reported in the common bean (*Phaseolus vulgaris*) (Bellucci et al., 2014) and subsequently in other domesticated plants and animals (Sauvage et al., 2017; Liu et al., 2019; Burgarella et al., 2021). Similar observations have been reported at the level of metabolic diversity (Beleggia et al., 2016; Alseekh et al., 2021).

Changes in nucleotide and gene expression diversity during the domestication of tetraploid wheat (*Triticum turgidum* L., 2n = 4x = 28; AABB genome) are not fully understood. Evidence indicates that domestication occurred in two well-defined phases: Primary domestication from wild emmer (*Triticum turgidum* ssp. *dicoccoides*) to emmer (*Triticum turgidum* ssp. *dicoccum*) started ∼12,000 years ago in the Fertile Crescent. This was followed by secondary domestication from emmer to durum wheat (*Triticum turgidum* ssp. *durum*), which started ∼2,000 years ago in the Near East and gave rise to durum wheat, the most important form of tetraploid wheat and currently the most widespread Mediterranean crop (Gioia et al., 2015; Taranto et al., 2020).

The molecular mechanisms underlying phenotypic plasticity in crops (Laitinen and Nikoloski, 2019) and their wild relatives must be understood to address the challenges faced by modern agriculture, including the overreliance on nitrogen (N) fertilizers to meet Sustainable Development Goals (SDGs). N is an essential macronutrient whose availability is directly linked to crop yield and grain quality (protein content) (Barneix, 2007; Howarth et al., 2008; Laidò et al., 2013), but it is also harmful to people and nature. Indeed, excess of N from agricultural sources is one of the major pollutant in fresh water (Bijay-Singh and Craswell, 2021). Understanding genetic variations in N acquisition, assimilation and metabolism can therefore provide novel sustainable strategies for crop improvement (Plett et al., 2018; Hawkesford and Griffiths, 2019). In tetraploid wheat, phenotypic differences related to N availability primarily arose during secondary domestication rather than primary domestication (Gioia et al., 2015), but the relationship between N metabolism and changes in gene expression plasticity during domestication is unclear.

Here we analysed 32 wild emmer, emmer and durum wheat genotypes by RNA-Seq to determine how contrasting differences in N availability shaped the nucleotide and gene expression diversity of tetraploid wheat during primary and secondary domestication. Our results provide insight into the pivotal role of N during the domestication and adaptive plasticity of one of our major food crops.

## Results and discussion

### A greater loss of nucleotide diversity occurred during the secondary domestication of tetraploid wheat

We prepared 128 RNA-Seq libraries from 4-week-old leaves of 32 tetraploid wheat genotypes representing *T. turgidum* ssp. *dicoccoides*, ssp. *dicoccum* and ssp. *durum* (Supplementary Table S1). On average, 6.8 million of reads per genotype (Supplementary Table S1) were mapped to the A and B reference subgenomes of bread wheat (Alaux et al., 2018). The mapping frequency exceeded 85% for all the three subspecies and the fraction of reads mapping to gene regions exceeded 72% (Supplementary Table S1).

Variant calling produced 800,996 high-quality single-nucleotide polymorphisms (SNPs). The number of polymorphic sites was similar in wild emmer (617,128) and emmer (613,509), but was much lower in durum wheat (425,513), confirming the higher genetic diversity of the wild population. We identified 190,377 common SNPs shared by all three taxa. As expected, wild emmer and emmer shared the highest percentage of SNPs (33%, 206,578). In contrast, durum wheat shared only 11% (46,352) of its SNPs with wild emmer and 17% (71,147) with emmer.

SNPs principal component analysis (PCA) revealed the broad genetic structure of the three wheat taxa (Figure 1) and confirmed that secondary domestication had a greater impact than primary domestication in differentiating the durum wheat subspecies. The analysed 12 durum wheat genotypes are genetically very similar, forming a dense cluster that is clearly distinguishable from the wild emmer and emmer genotypes. In contrast, the wild emmer and emmer genotypes were loosely clustered, indicating a greater genetic admixture. These results are consistent with previous genetic studies on the origins of domesticated wheat and reflect the multiple stages of domestication (Luo et al., 2007; Civáň et al., 2013; Oliveira et al., 2020), and indicate that the used genotypes are representative.

**Figure 1:**
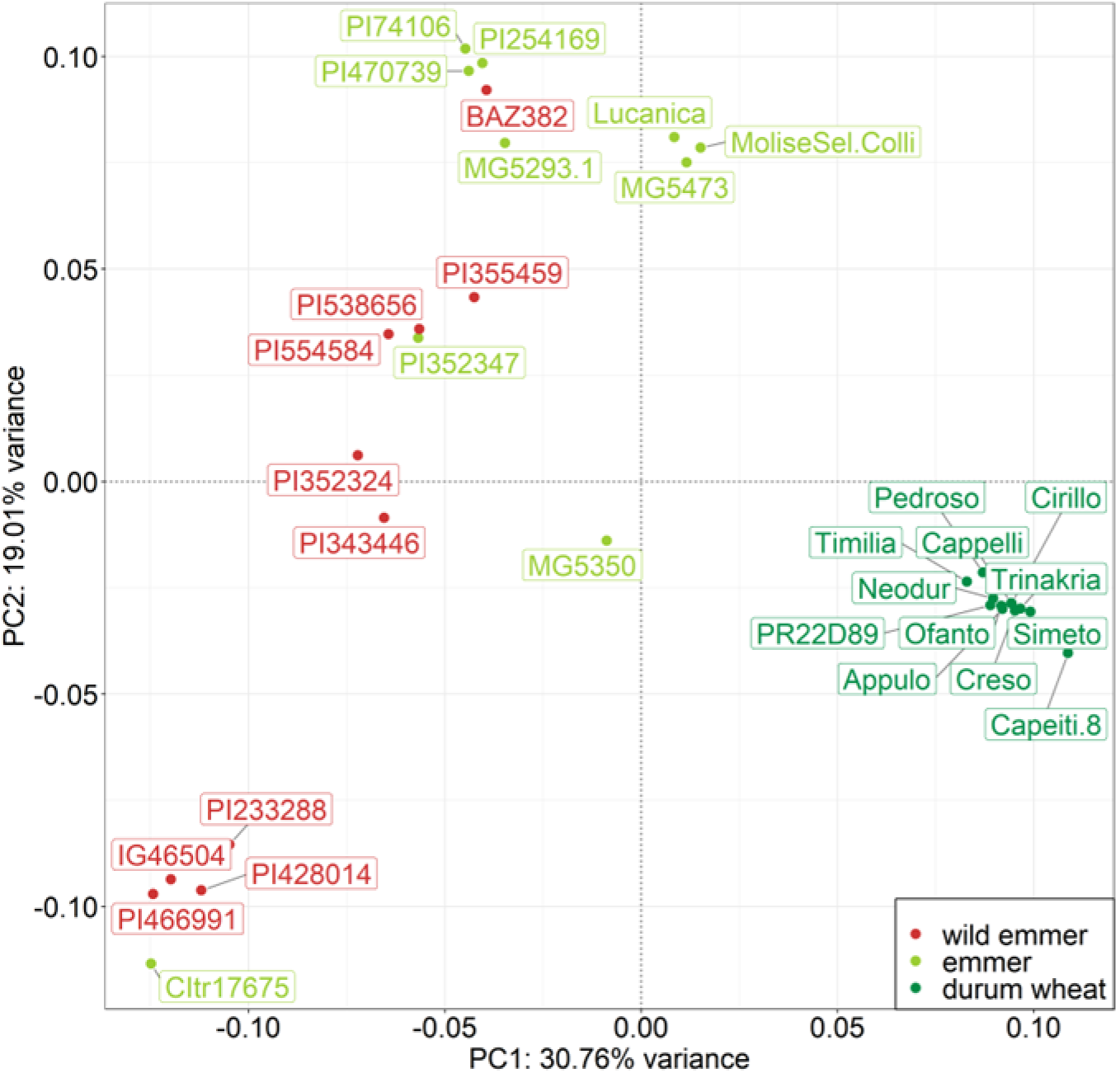
Principal component analysis of 32 wheat genotypes based on single-nucleotide polymorphisms (SNPs). The first two principal components (PC1 and PC2) are shown. The three colors represent different taxa. Labels show the accession name of each genotype.

Nucleotide diversity estimates (Table 1) show the expected substantial loss of nucleotide diversity during domestication. The average nucleotide diversity of durum wheat was ∼35% lower than domesticated emmer, which was in turn ∼11% lower than wild emmer, highlighting the greater impact of secondary domestication. When the cumulative effect of primary and secondary domestication is taken into account, we observed a ∼42% reduction in the nucleotide diversity of durum wheat compared to its wild ancestor (Table 1).

**Table 1:**
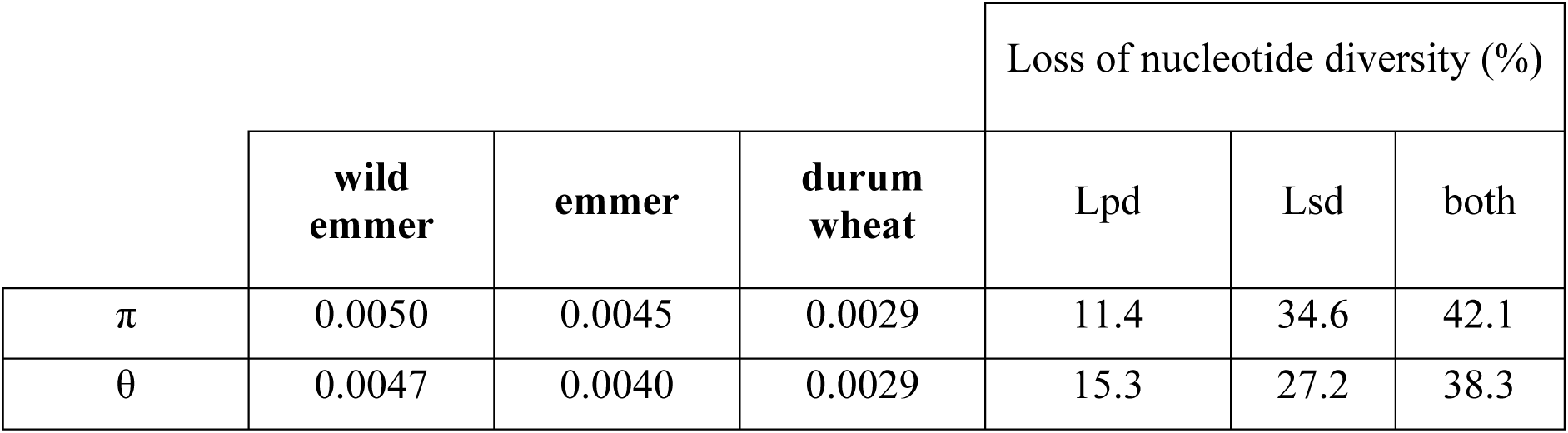
Nucleotide diversity estimates and diversity loss for the three wheat taxa. Diversity loss is shown during primary domestication (wild emmer to emmer, Lpd), secondary domestication (emmer to durum wheat, Lsd) and both processes (wild emmer to durum wheat), based on average π and θ values. The π and θ symbols represent averaged estimates of nucleotide diversity.

### The variability of gene expression during domestication was influenced by N availability

To quantify the diversity of gene expression in each subspecies, we calculated evolvability scores under high and low N availability conditions. Evolvability was estimated using the additive coefficient of variation (CV_A_) in read counts (Supplementary Table S2). In contrast to heritability, CV_A_ is a standardized measure of additive genetic variation that is not influenced by other sources of variance (Houle, 1992; Hansen et al., 2011), and is therefore well suited for comparative analysis (Garcia-Gonzalez et al., 2012). As for nucleotide diversity, we found that the CV_A_ decreased during domestication under both N conditions; however, the mean CV_A_ of all three subspecies was higher under low N conditions (Figure 2a,b; Table 2). High N availability therefore appears to promote a more uniform gene expression pattern, whereas higher variability is observed during N starvation. The association between domestication and declining diversity in gene expression has also been reported in crops, such as: common bean (Bellucci et al., 2014), tomato (Sauvage et al., 2017) and sorghum (Burgarella et al., 2021) as well as domesticated animal species (Liu et al., 2019).

**Figure 2:**
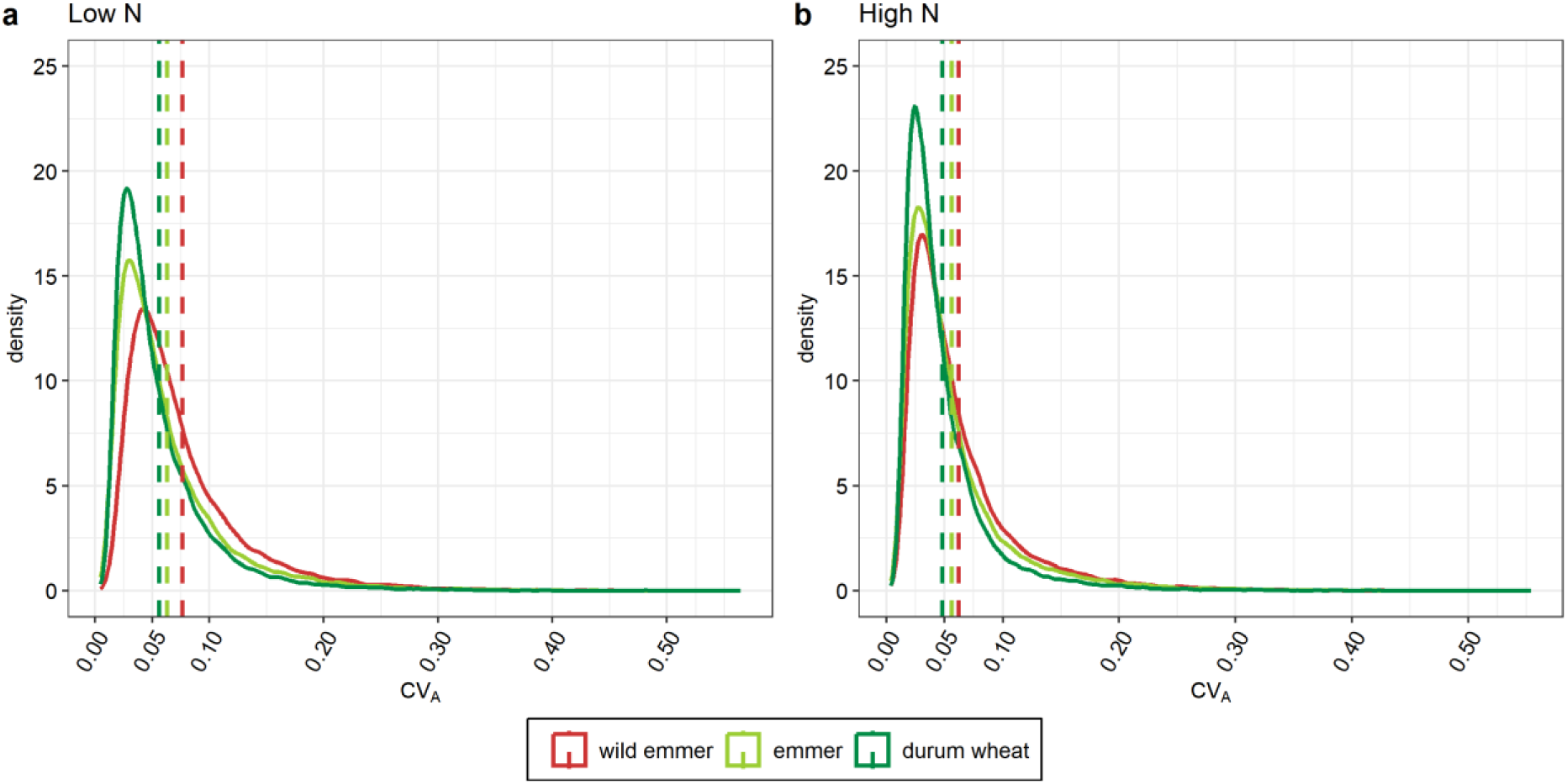
Density plots of the additive coefficient of variation (CVA) in the three wheat taxa. Comparison of the estimated density functions of the CV_A_ in gene expression, calculated using all 32,358 genes. **a** High nitrogen conditions. **b** Low nitrogen conditions. Dashed lines represent the averaged CV_A_ value, colored according to the different taxa.

We used the contrasting N conditions of our samples to examine whether the loss of expression diversity is associated with the specific aspects of the cultivation environment, causing primary and secondary domestication to have a significantly different impact. Under high N conditions, we observed a ∼9% loss in expression diversity in emmer compared to wild emmer (effect of primary domestication) and a ∼15% loss in durum wheat compared to emmer (effect of secondary domestication). In contrast, these losses were ∼18% and 11% under N starvation conditions, revealing twice the loss of expression diversity during primary domestication, but a lower value during secondary domestication (Table 2). All four values differed significantly from each other (Mann–Whitney test, p < 0.001). The opposing expression diversity profiles during domestication under high and low N conditions were observed not only for overall gene expression, but also for the subgroup comprising all differentially expressed genes (DEGs) and the subgroup comprising all unmodulated genes (Supplementary Table S3). The loss of expression diversity among the DEGs due to primary domestication was ∼9% and ∼15% under high and low N conditions, respectively, whereas the loss due to secondary domestication was ∼18% and ∼14% under high and low N conditions, respectively (Supplementary Table S3). The loss of expression diversity among the unmodulated genes was similar to the values for overall gene expression (Supplementary Table S3).

**Table 2:**
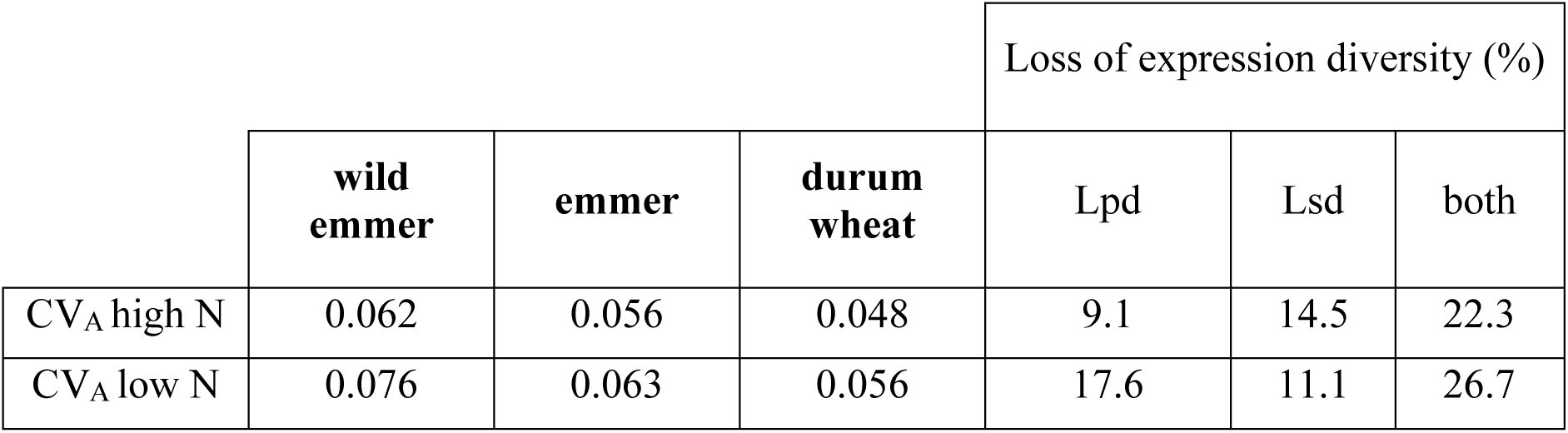
Mean additive coefficient of variation (CVA) in gene expression and loss of expression diversity for the three wheat taxa. Diversity loss is shown during primary domestication (wild emmer to emmer, Lpd), secondary domestication (emmer to durum wheat, Lsd) and both processes (wild emmer to durum wheat), based on averaged CVA values calculated for all 32,358 genes.

A phenotypic study of the same accessions used in the present work has already shown that secondary domestication reduced the phenotypic diversity under high N conditions, but the reduction was smaller and not significant under N starvation conditions (Gioia et al., 2015). In the case of durum wheat, selection has apparently enhanced the growth response to N availability, indicating a putative focus on improving N uptake and utilization efficiency. Our expression diversity results indicate that selection has favored specific traits and thus led to a more uniform set of cultivars, as also suggested in earlier study using morphological traits (Gioia et al., 2015).

### Domestication and nitrogen availability shaped the divergence of tetraploid wheats

Genetic differentiation among the three subspecies was estimated by calculating the pairwise fixation index (*F_ST_*) for every gene locus in our dataset. As shown in Figure 3a, the lowest genetic differentiation was observed between wild emmer and emmer (mean *F_ST_* = 0.09), whereas much higher genetic differentiation was found between emmer and durum wheat (mean *F_ST_* = 0.27) and, similarly, between wild emmer and durum wheat (mean *F_ST_* = 0.28). These values align with earlier findings that examined broad collections of tetraploid wheat accessions (Luo et al., 2007; Mazzucotelli et al., 2020), and provide additional evidence for the representativeness of the genotypes used.

**Figure 3:**
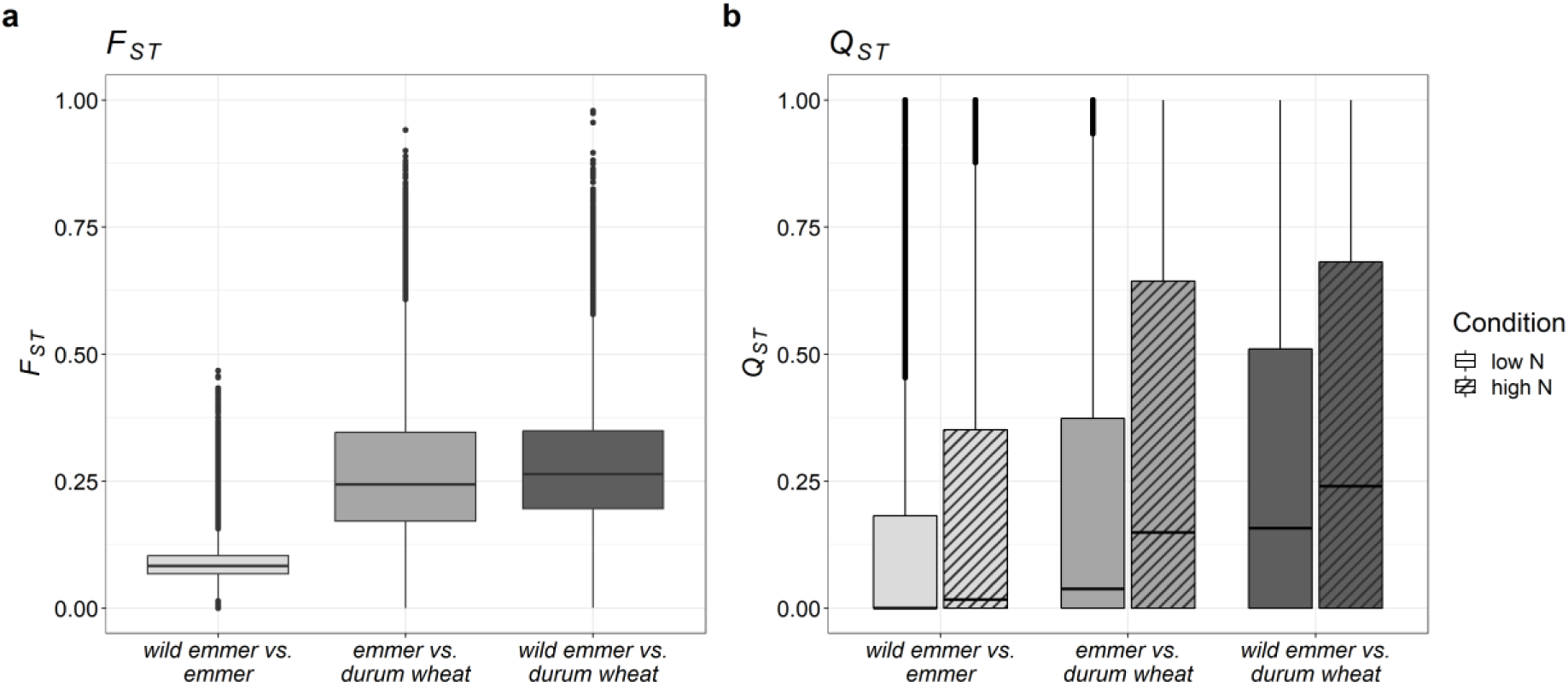
***F****_ST_* **and *Q****_ST_* **distributions. a** Boxplots showing the gene locus *F_ST_* distribution for every subspecies pairwise comparison. **b** Boxplots showing the transcript *Q_ST_* distribution for every subspecies pairwise comparison under low nitrogen and high nitrogen conditions, represented by empty and hatched grayscale bars, respectively.

Divergence at the transcriptomic level was estimated by calculating *Q_ST_*, the quantitative analog of *F_ST_*, taking N availability into account as an environmental variable. Under both N conditions, we observed the same trend shown for *F_ST_* (Figure 3b). Specifically, secondary domestication had a stronger impact on differentiation (emmer *vs* durum wheat, mean *Q_ST LN_* = 0.23, mean *Q_ST HN_* = 0.33) than primary domestication (wild emmer *vs* emmer, mean *Q_ST LN_* = 0.16, mean *Q_ST HN_* = 0.23). Interestingly, the *Q_ST_* distributions of every pairwise comparison showed higher values under high N conditions compared to N starvation (Figure 3b), suggesting that N availability during domestication significantly contributed to the differentiation of gene expression in tetraplid wheats.

The *Qst* distributions were used to perform a “selection scan”, seeking genes whose expression was potentially under selection. Starting from 5,868 genes meeting the heritability criteria (*H^2^* ≥ 0.7 or *S×N* ≥ 0.2, that is the species × environment variance component i.e., every species subgroup × N condition; Supplementary Figure S1), we retained 973 genes having *Qst* values in the 5% right tail of the distributions. The *Qst–Fst* comparison method (Leinonen et al., 2013) was then used to confirm that the divergent expression (high *Qst* values) of the filtered genes was caused by directional selection (*Qst* > *Fst*) and not by genetic drift (*Qst* ≈ *Fst*) or stabilizing selection (*Qst* < *Fst*) (Leinonen et al., 2013). After removing *Fst* values < 0.01, we retained 967 genes satisfying the criterion *Q_ST_* > *F_ST_*, indicating that their expression was likely subjected to directional selection in at least one of the evolutionary contexts examined herein (i.e., primary and/or secondary domestication under high and/or low N availability conditions) (Supplementary Table S4).

Gene Ontology (GO) enrichment analysis revealed that selection acted on distinct gene categories during primary and secondary domestication (Supplementary Figure S2). During primary domestication, we found categories associated with “defense-related programmed cell death, modulated by biotic interactions”, indicating an enhanced plant hypersensitive response to pathogens. This can be interpreted as a consequence of the transition from the natural growing environment of the wild genotypes to agroecosystems characterized by high-density domesticated crop monocultures. In this context, crops face higher disease pressure from crop-specific pathogens (Savary et al., 2019) and therefore induce a hypersensitive response, which can lead to programmed cell death and necrosis as a defense mechanism. It is important to note that pathogen defense mechanisms in plants often overlap with the regulation of beneficial symbiotic interactions, therefore, one expects a trade-off between traits associated with symbiosis and innate immunity (Porter and Sachs, 2020). Moreover, domesticated crops are less able to fully benefit from microbial interactions than their wild relatives, as observed in a comparative study of bread wheat landraces as well as old and modern varieties (Valente et al., 2023). One contributing factor is the widespread use of high-input agricultural practices, because the availability of fertilizers reduces the need for plants to invest in symbiotic relationships (Martín-Robles et al., 2018). Additionally, certain target traits in plant breeding, such as phytohormones that regulate flowering time and plant height, can have unintended effects on beneficial symbiosis due to pleiotropy (Sawers et al., 2018).

Among the genes found to be under selection during secondary domestication, we observed the enrichment of categories associated with amino acid metabolism, particularly those related to the “lysine catabolic process” (Supplementary Figure S2). This included genes encoding the bifunctional enzyme lysine ketoglutarate reductase/saccharopine dehydrogenase (LKR/SDH). This enzyme is ubiquitous in plants and animals, and represents the key step in lysine catabolism via the saccharopine pathway (SACPATH). The structure and transcription of the *LKR/SDH* gene has been studied in *T. durum* and compared with other plants, showing species-dependent differences in expression levels including lineage-specific differences between monocots and dicots (Anderson et al., 2010). Lysine is the first limiting essential amino acid in cereal grains and its catabolic pathway has been targeted to increase the lysine content of maize and rice seed (Houmard et al., 2007; Frizzi et al., 2008; Long et al., 2013). Generally, the quantity of lysine-containing proteins in cereal seeds is much lower than that of storage proteins devoid of lysine, such as prolamins (specifically gliadin in wheat). The SACPATH seems to channel the lysine skeleton into the production of glutamic acid, which is a precursor of proline, one of the most abundant amino acids in glutens (Arruda et al., 2000).

General changes in amino acid metabolism during domestication have been observed in other crops based on nucleotide data, including sunflower (Chapman et al., 2008), maize (Swanson-Wagner et al., 2012) and common bean (Bellucci et al., 2014). Evolutionary metabolomics has also revealed signatures of selection affecting amino acid metabolism during secondary domestication (Beleggia et al., 2016). In durum wheat, domestication was linked to the selection of a specific protein composition and led to a notable decrease in the diversity of gliadin and glutenin subunits, strongly correlating with grain yield and the technological properties of gluten (Laidò et al., 2013). The analysis of spring wheat genotypes has shown that the SACPATH is upregulated in response to drought stress, and is significantly more active in drought-tolerant compared to drought-susceptible genotypes (Michaletti et al., 2018). This may reflect the role of proline, which can be produced from this pathway, as a major constituent of storage proteins and one of the main osmoprotectants produced as a response to stress (Kavi Kishor et al., 2022). These findings suggest that selection for stress-tolerant genotypes as well as seed protein composition during wheat domestication influenced the expression of SACPATH genes.

### Changes in nitrogen availability trigger gene expression, resulting in a twofold increase in the number of differentially expressed genes in durum wheat compared to emmer and wild emmer wheat

We identified DEGs in each subspecies that discriminated between high N conditions and N starvation using a stringent pipeline and strict thresholds (p-adjust < 0.001) to reduce the number of false positives. We found 3,326 DEGs in wild emmer, 3,305 in emmer and 5,901 in durum wheat, with more upregulated than downregulated genes in all three subspecies. Durum wheat had the highest percentage of private DEGs (∼42%, 2,479), whereas similar numbers were found in wild emmer (∼14%, 458) and emmer (∼15%, 486). Wild emmer and emmer shared ∼23% (749) and ∼21% (700), respectively, of their DEGs with durum wheat. The percentage of DEGs shared only between wild emmer and emmer was 4% (146), but almost 60% of wild emmer and emmer DEGs and ∼33% of durum wheat DEGs were shared by all three taxa (Figure 4a). The proportions of private and shared DEGs were preserved when we separated them into upregulated and downregulated subsets (Figure 4b,c). In all three taxa, most DEGs were located on chromosomes 2A, 2B, 3A, 3B, 5A and 5B, each carrying > 7.5% of the DEGs; in contrast, chromosomes 6A and 6B each contained only ∼5% of the DEGs Supplementary Figure S3.

**Figure 4:**
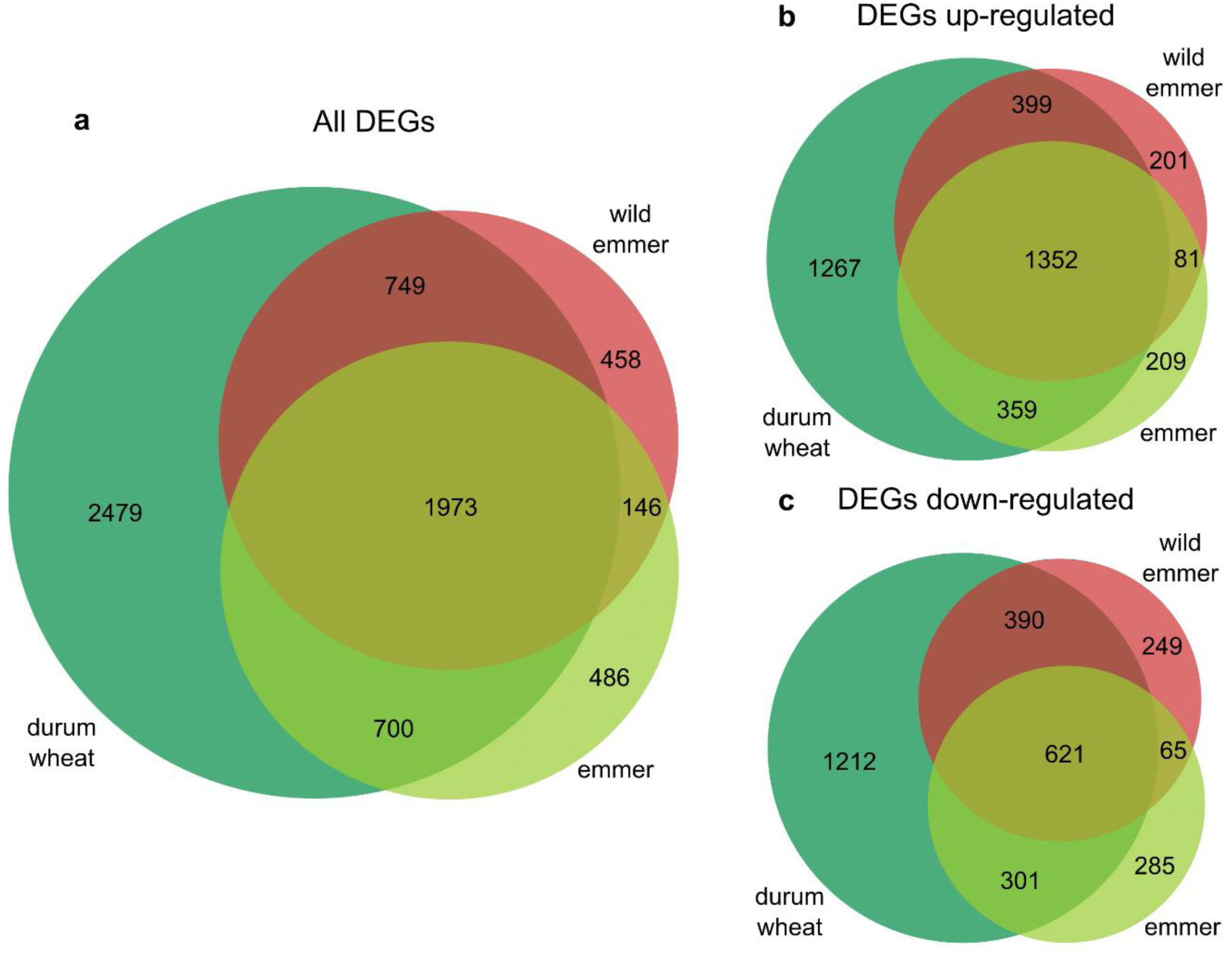
Differentially expressed genes (DEGs) when comparing high and low nitrogen conditions within each subspecies. Venn diagrams showing **a** Total set of DEGs; **b** upregulated DEGs only; and **c** downregulated DEGs only.

GO enrichment analysis of the DEGs meeting the threshold FDR < 0.05 revealed 23 macro-categories in wild emmer, 21 in emmer and 25 in durum wheat (Supplementary Figure S4). The main differences between the three subspecies were observed for categories related to “signaling”, “regulation of biological process”, “developmental process”, and “metabolic process” Supplementary Table S5. We observed the uniform enrichment of GO categories associated with upregulated genes in all three subspecies, including terms linked to N and amino acid metabolism as well as carbon metabolism and photosynthesis (Supplementary Table S5). In contrast, the enrichment of GO categories associated with downregulated genes was more selective, with some GO categories related to N metabolism enriched only in durum wheat, including GO:0006807 and GO:0034641 (N compound and cellular N compound metabolic process, respectively) and GO:0006536 “glutamate metabolic process” (Supplementary Table S5). Functional annotations of the most strongly modulated genes (top 5% |log_2_FC| values) are reported in Supplementary Table S6.

Our data confirm, on a larger set of samples, earlier observations on the response of wheat to N starvation based on transcriptomics and metabolomics data^36–38^. These earlier studies included one emmer and one durum wheat genotype also present in our sample set (Beleggia et al., 2021), but also considered the durum wheat cultivar Svevo (Curci et al., 2017) and various bread wheat cultivars (Sultana et al., 2020). As expected, genes involved in N metabolism were modulated during N starvation. Among the key genes for N assimilation, those encoding asparagine synthetase and nitrite reductase were upregulated in every taxon, whereas those encoding glutamate carboxypeptidase and glutamate decarboxylase were downregulated. We observed contrasting profiles for genes encoding ureide permease (encoding a ureide transporter), which were strongly upregulated in all three subspecies in response to N stress, whereas genes encoding nitrate transporters were strongly downregulated. The modulated genes also included transporters of amino acids and other nutrients.

N starvation also influenced other metabolic pathways, revealing many further DEGs involved in C metabolism, especially fatty acid metabolism, glycolysis, photosynthesis and the tricarboxylic acid (TCA) cycle. About 10% of the highest-ranking DEGs represented transcription factors and protein kinases. The most common functional category (accounting for 17% of annotated DEGs) reflected the general stress response to N starvation, including the mitigation of oxidative stress and detoxification. Examples included genes encoding *cytochrome P450s*, *glutaredoxin family*, *glutathione S-transferases* and *peroxidases* (Supplementary Table S6).

To compare gene expression between the three taxa while taking the environmental effects into account, we also identified DEGs between each pair of subspecies under all N conditions. Accordingly, we compared emmer *vs* wild emmer (primary domestication, high and low N), durum wheat *vs* emmer (secondary domestication, high and low N) and durum wheat *vs* wild emmer (cumulative effect, high and low N) (Supplementary Figure S5). The wild emmer *vs* emmer comparison revealed few DEGs regardless of N availability (12 and 11 DEGs under high and low N conditions, respectively), whereas the emmer *vs* durum wheat comparison revealed 41 DEGs associated with high N and 29 associated with N starvation, and the wild emmer *vs* durum wheat comparison revealed 46 DEGs associated with high N and only 10 associated with N starvation. These data indicate that the number of DEGs increases during domestication but only when there is a sufficient N supply (Supplementary Figure S5). Interestingly, there were more upregulated than downregulated genes in all pairwise comparisons under high N conditions (∼65%) but the proportion increased under N starvation conditions, particularly for the comparison of wild emmer *vs* durum wheat (90%). The preponderance of upregulated genes during domestication has also been observed in maize (Lemmon et al., 2014), whereas domestication was shown to increase the proportion of downregulated genes in common bean (Bellucci et al., 2014), egg-plant (Page et al., 2019) and sorghum (Burgarella et al., 2021) landraces compared to wild relatives. The absence of consistent patterns suggests that the evolution of domesticated phenotypes is driven by specific processes that are unique to each crop.

Among the 102 DEGs (Supplementary Table S7) found in at least one of the six pairwise comparisons between subspecies, 35 were also found among DEGs identified between contrasting N conditions and of which 24 were proposed to be under selection. Overall, six genes were identified in all three experiments (i.e., differentially expressed between subspecies and between contrasting N conditions, and showed evidence of selection).

### Selection shaped the expression profiles of genes modulated by nitrogen availability

The 6,991 DEGs found in at least one species when comparing the contrasting N conditions included 101 putatively under selection, which are candidates for the adaptive response to N availability. We applied PCA to the normalized read counts in order to investigate if the different genotype groups can be separated based on their gene expression. Initially we incorporated all 6,991 DEGs (Figure 5a,b) before focusing on the subset of 101 DEGs that were also putatively under selection (Figure 5c,d). When considering all DEGs, PC1 did not completely separate the durum wheat genotypes from the other taxa, in contrast to the clear separation observed for the SNP data (Figure 1), and this was particularly evident during N starvation (Figure 5b). There was also a moderate degree of overlap between the wild emmer and emmer genotypes along PC2. However, when we focused on the DEGs under selection, PC1 separated the durum wheat genotypes into a densely clustered group (as observed for the SNP data) under both N conditions, and PC2 separated the wild emmer and emmer genotypes more clearly, especially under high N conditions (Figure 5c,d).

**Figure 5:**
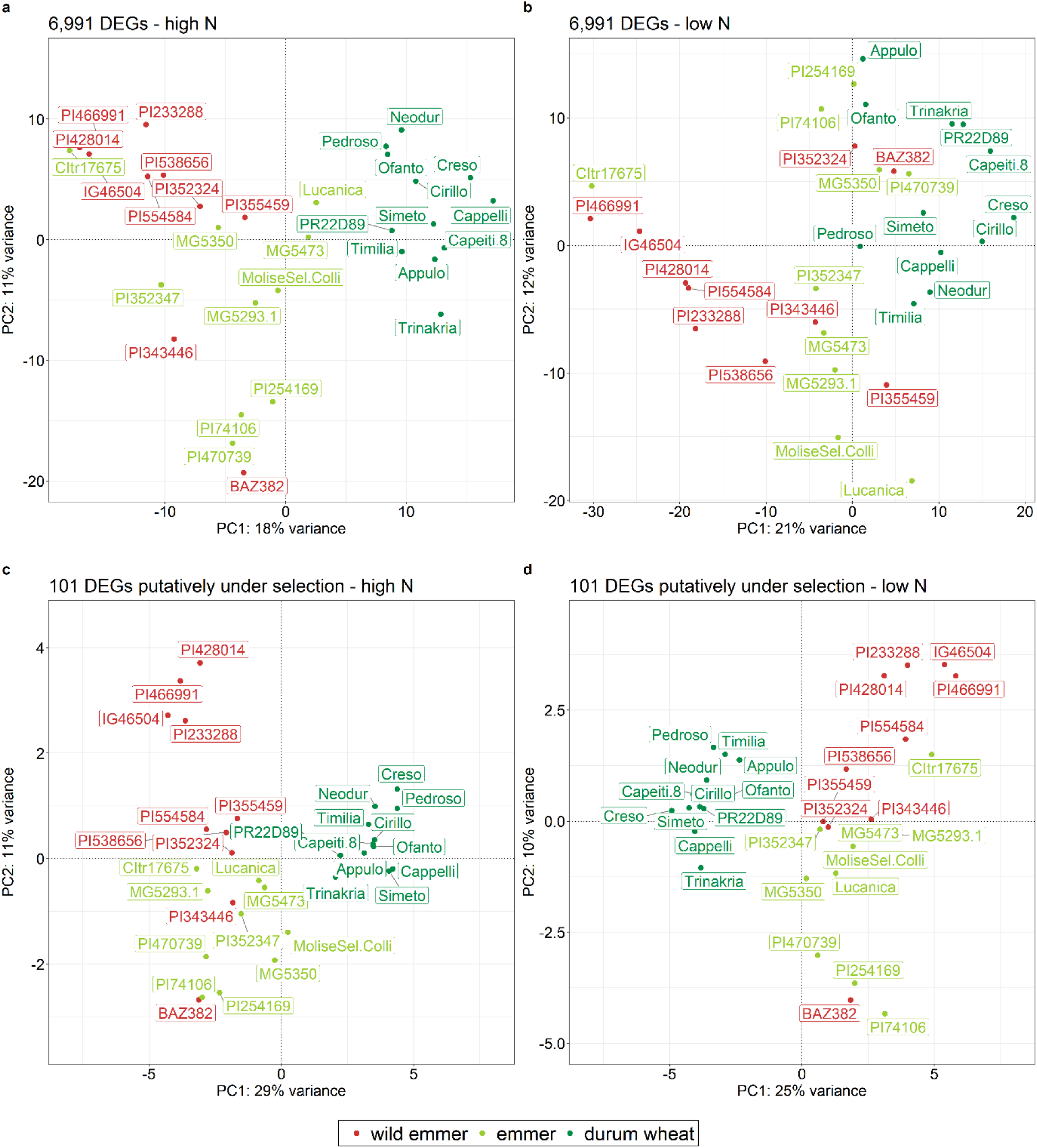
Principal component analysis of differentially expressed genes when comparing high and low nitrogen conditions within each subspecies. a,b. Plots based on all 6,991 DEGs (not filtered): **a** high nitrogen conditions and **b** low nitrogen conditions. **c,d** Plots based on 101 DEGs that are also putatively under selection: **c** high nitrogen conditions and **d** low nitrogen conditions. Samples are represented by taxa-based colored dots. Labels show the accession name of each genotype.

The integration of selection signatures (based on *Q_ST_/F_ST_* values) and differential expression analysis uncovered a set of 101 candidate genes that are interesting due to their potential roles in the domestication and diversification of cultivated wheat, specifically in relation to N availability. Functional annotation (Supplementary Table S8) revealed upregulated genes involved in carbon (C) metabolism as well as some transcription factors and transporters, as well as both upregulated and downregulated genes responsible for general stress responses and N metabolism, specifically those encoding enzymes involved in amino acid metabolism, such as methionine aminopeptidase, aspartokinase and glutamate dehydrogenase (GDH). The latter is particularly noteworthy because, in addition to its modulation in response to different N conditions and the presence of selection signatures, it was also upregulated in the comparison between wild emmer and durum wheat under high N conditions. GDH is a key enzyme involved in N metabolism and N/C balance (Miflin and Habash, 2002). This is supported by the co-localization of quantitative trait loci for GDH activity and physiological traits associated with the flag leaf lamina, such as soluble protein and amino acid content, as well as flag leaf area and dry weight (Fontaine et al., 2009). Selection signatures were also identified in the *GDH* gene when comparing landraces with old and modern durum wheat cultivars (Taranto et al., 2020). Our results confirm that N metabolism has been a key driver during the evolutionary history of wheat, particularly the central role of glutamate in the process of domestication. This was also suggested by a combined transcriptomics and metabolomics study, showing that glutamate and γ-aminobutyric acid (mainly synthetized from glutamate) are central to the genotype-specific response of emmer and durum wheat to N starvation (Beleggia et al., 2021).

We have shown that significant changes occurred at the nucleotide and gene expression levels during the domestication of tetraploid wheat, taking into account the environmental variable of N availability. We confirmed that more nucleotide diversity has been lost during secondary domestication compared to primary domestication, and revealed a parallel trend in the loss of gene expression diversity associated to the domestication process, with a stronger effect due to secondary domestication and unveil a parallel different impact of primary and secondary domestication on the loss of expression diversity, which appears to be related to N availability in the durum wheat selection environment. We present evidence that selection may have operated in different directions during primary and secondary domestication, the former involving changes related to biotic interactions and the latter related to amino acid metabolism. By screening a large number of genotypes, we found a major transcriptional response in durum wheat (compared to emmer and wild emmer) to changes in N availability. Finally, through the innovative combination of RNA-Seq analysis and the estimate of quantitative genetics parameters, we developed a pipeline to identify selection signatures and phenotypic plasticity in gene expression data based on evolvability and *Q_ST_/F_ST_* scores. Our findings, elucidating the role of N in tetraploid wheat domestication and adaptive response can guide the development of innovative strategies for crop improvement, resource use efficiency, and environmental sustainability.

## Materials and methods

### Plant material and experimental design

The study included 32 tetraploid wheat genotypes, comprising 10 accessions of wild emmer (*T. turgidum* ssp. *dicoccoides*), 10 accessions of emmer (*T. turgidum* ssp. *dicoccum*), and 12 accessions of durum wheat (*T. turgidum* ssp. *durum*) (Supplementary Table S1). The samples we analysed were part of a larger experiment, conducted in October 2012 and described elsewhere (Gioia et al., 2015). Briefly, wheat genotypes were grown for 4 weeks under high nitrogen (N+) and nitrogen starvation (N–) conditions in the Phytec Experimental Greenhouse at the Institute of Biosciences and Geosciences (IBG-2), Plant Sciences Institute, Forschungszentrum Jülich GmbH, Germany (50°54′36′′ N, 06°24′49′′ E). Seeds of uniform size and mass were visually selected, surface sterilized (1% (*w/v*) NaClO for 15 min) and pre-germinated. After germination, seedlings showing uniform growth (seminal root length, 1–2 cm) were transferred to soil-filled rhizoboxes, which were placed in the automated GROWSCREEN-Rhizo phenotyping system available at IBG-2. We used a Type 0 manually sieved peat soil (Nullerde Einheitserde; Balster Einheitserdewerk, Frondenberg, Germany), which provided low nutrient availability (ammonium N and nitrate N concentrations of < 1.0 and < 1.0 mg l^−1^, respectively). All plants were watered twice daily with 400 ml of tap water and were supplied three times per week with 200 ml of modified Hoagland solution(Hoagland and Arnon, 1950) with or without added N. For the N starvation conditions, KNO_3_ and Ca(NO_3_)_2_ were replaced with K_2_SO_4_ and CaCl_2_·6(H_2_O), respectively. The experiment was carried out under natural lighting in the greenhouse, with an air temperature of 18–24 °C and a relative humidity of 40–60%. For each N treatment, we used two replicates of each genotype with two plants per replicate (four plants per genotype in total). After 4 weeks, leaves were pooled from two plants of the same genotype growing in the same rhizobox. Accordingly, four independent biological replicates (two replicates per N condition) were produced for each genotype, with the exception of wild emmer IG 46504, PI 233288, PI 466991, PI 538656, emmer MG 5293/1, and durum wheat Creso, Pedroso and Trinakria, for which only three replicates were available, and emmer Molise Sel. Colli and durum wheat Simeto, for which eight replicates were available. The tissues were immediately frozen in liquid N_2_ and stored at −80 °C. Further details of the experiment and growth conditions are provided elsewhere(Gioia et al., 2015).

### RNA extraction and sequencing

RNA was extracted from 100 mg of frozen ground leaves per replicate using the Spectrum Plant Total RNA kit (Sigma-Aldrich, St Louis, MO, USA) followed by treatment with RNase-free DNase using the On-Column DNase I Digestion Set (Sigma-Aldrich). RNA integrity and purity were assessed by agarose gel electrophoresis and a Bioanalyzer 2100, respectively (Agilent/Bonsai Technologies, Santa Clara, CA, USA). Only RNA samples with an RNA integrity number > 8.0 were considered suitable for analysis.

Library construction and RNA sequencing were carried out using the Illumina mRNA-Seq platform at the Montpellier Genomix sequencing facility (http://www.mgx.cnrs.fr) as previously described (David et al., 2014). Briefly, RNA samples were processed using TruSeq RNA sample preparation kits v2 (Illumina, San Diego, CA, USA). Libraries were quantified by real-time PCR using the KAPA Library Quantification Kit for Illumina Sequencing Platforms (Roche, Basel, Switzerland), followed by quality control using a DNA 100 Chip on a Bioanalyzer 2100. Cluster generation and sequencing were carried out using the Illumina HiSeq 2000 instrument and TruSeq PE Cluster Kit v3, following the Illumina PE_Amp_Lin_Block_V8.0 recipe, and Illumina TruSeq PE Cluster v3-cBot-HS kits with the 2 × 100 cycles, paired-end, indexed protocol, respectively (David et al., 2014).

### RNA-Seq library processing and mapping

We pre-processed 128 raw paired-end RNA-Seq libraries (David et al., 2014). Cutadapt (Martin, 2011) was then used to remove adaptor sequences and trim the end of reads with low quality scores (parameter –q 20) while keeping reads with a minimum length of 35 bp. Reads with a mean quality score < 30 were discarded, and orphan reads (whose mates were discarded in the previous filtering steps) were removed (David et al., 2014). The final quality of trimmed and filtered reads was assessed using FastQC (Andrews, 2014).

The bread wheat (*Triticum aestivum* cv. Chinese Spring) genome assembly IWGSC RefSeq v2.1, along with the corresponding genome annotation, were downloaded from the IWGSC data repository hosted by URGI-INRAE (https://wheat-urgi.versailles.inra.fr/) and used as a reference to map each cleaned library to the A and B sub-genomes. The bread wheat genome was chosen deliberately to ensure the inclusion of an outgroup species that is closely related to the subspecies in the panel. By doing so, we aimed to avoid bias that could arise from selecting only one subspecies among our panel of accessions. We have confidence in this strategy because the *T. aestivum* A and B subgenomes are derived from the tetraploid species included in the study.

STAR v2.7.0e (Dobin et al., 2013) was used for read mapping with the –-quantMode TranscriptomeSAM and –-quantTranscriptomeBan Singleend options. The output alignments were translated into transcript coordinates (in addition to alignments in genomic coordinates), allowing insertions, deletions and soft-clips in the transcriptomic alignments. The transcriptomic alignments were used as inputs for salmon v1.6.0 (Patro et al., 2017) to quantify gene expression. Raw read counts were computed for all genes in each sample and, to filter out weakly-expressed transcripts, only genes with at least 1 count per million (CPM) in at least 10 samples (of the same subspecies) were retained. This was calculated separately in each of the three subspecies and the raw counts of the filtered genes in each subspecies were then combined for downstream analysis, for a total of 32,358 genes (Supplementary Table S2).

### Variant identification

Variants were called by applying BCFtools v1.15 (previously SAMtools) (Danecek et al., 2021) to the alignment bam files. The “*bcftools mpileup*” command was used to determine the genotype likelihoods at each genomic position, with a minimum alignment quality of 20 and a minimum base quality of 30. The actual calls were obtained using the “*bcftools call*” command. The resulting VCF file was filtered using the “*bcftools view*” command, removing indels and keeping only sites covered by at least three reads in all genotypes. Subsequently, only biallelic SNPs with maximum values of 50% missingness and a 1% minor allele frequency were retained. To identify private and shared SNPs among the different subspecies, every possible comparison of the three subsampled VCF files (wild emmer, emmer and durum wheat) was carried out using the “*bcftools isec*” command.

### Population genetics analysis

Variants were filtered (one SNP per 500 kb) using the VCFtools *--thin 500000* option (v0.1.17) (Danecek et al., 2011) and then converted into ped format with PLINK (v1.90p) (Purcell et al., 2007). PLINK was also used to compute genetic distances between individuals with the *--distance-matrix* flag. The output matrix was used as input for PCA with the *cmdscale* function of R (v4.2.1) (R Core Team, 2022).

Genetic diversity statistics, including nucleotide diversity (π and θ) (Tajima, 1983; Watterson, 1975) were computed on the alignment bam files for each subspecies, from the folded site frequency spectra using ANGSD(Korneliussen et al., 2014). First, the *doSaf* function was used to estimate per-site allele frequencies (Saf) then *realSFS* was used to get the site frequency spectra. The statistical loss of diversity (Vigouroux et al., 2002) was used to test the impact of primary and secondary domestication on the molecular diversity of the three subspecies. For primary domestication, the statistic was computed as [1 − (x_emmer_/x_wild_)], where x_emmer_ and x_wild_ are the diversities in emmer and wild emmer, respectively, measured using π, θ and D. If x_emmer_ was higher than x_wild_, then the parameter was calculated as [(x_wild_/x_emmer_) – 1]. The loss of diversity due to secondary domestication in durum wheat versus emmer was calculated as [1 − (x_durum_/x_emmer_)], where x_durum_ and x_emmer_ are the diversities in durum wheat and emmer, respectively. If x_durum_ was higher than x_emmer_, then the parameter was calculated as [(x_emmer_/x_durum_) – 1].

We calculated *F_ST_* for each pair of populations using ANGSD (Korneliussen et al., 2014). Saf and 2D SFS were calculated as for nucleotide diversity, then the *fst index* function was used to obtain the global estimate. To get an *F_ST_* value for each gene in our dataset, we used the *fst print* function, which prints the posterior expectation of genetic variance between populations (called A), and total expected variance (called B) for every locus. We then computed the weighted *F_ST_* as the ratio of the summed As and summed Bs for every gene region, using an *ad hoc* R script.

### Expression profiles, heritability and *Q_ST_* analysis

Raw read counts of the 32,358 genes were normalized using the *vst* method allowing the additive coefficient of variation (CV_A_) (standard deviation/mean) to be calculated for the two N conditions in every subspecies, averaging the biological replicates of every genotype. The statistical loss approach (Vigouroux et al., 2002) was then applied to test the loss of expression diversity in the different groups, as previously reported (Bellucci et al., 2014). The statistical significance of the differences between each CV_A_ value and the percentage loss of expression diversity was determined using the Mann-Whitney test in R (v4.2.1) (R Core Team, 2022) with the function *wilcox.test*.

To compute heritability, the raw counts of each subspecies under each condition were first normalized using the trimmed mean M-values normalization method in the R package edgeR(Robinson et al., 2010) and the voom normalization method in the R package limma(Smyth, 2005). To determine the variance component of each factor and heritability, the following model was considered:

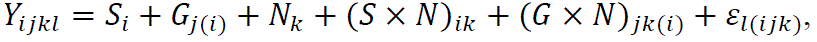

where *Y*_*ijkl*_ is the normalized gene expression level, *S*_*i*_ is the species factor, *G*_*j*(*i*)_is the genotype factor nested in species, N_*k*_ is the N level factor, (*S* × N)_*ik*_is the interaction between species and N levels, (*G* × N)_*jk*(*i*)_is the interaction between genotypes and N levels, and ε_*l*(*ijk*)_is the residual error. All factors were treated as random effects in the model except the intercept, which was a fixed effect. The linear mixed models were fitted using the *lmer* function in R package lme4 based on the normalized data of each transcript(Bates et al., 2015). The heritability (H^2^) was calculated as 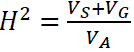, where 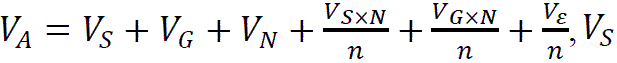 is the variance of species, *V*_*G*_ is the variance of genotype, *V*_N_is the variance of N level, *V*_*S*×N_is the variance of species and N level interaction, *V*_*G*×N_ is the variance of genotype and N level interaction, *V*_ε_ is the residual variance, and *n* is the number of N levels. *V*_*S*×N_ and *V*_*G*×N_ represent the genotype × environment interaction variance components at the species and genotype (nested in species) levels, respectively.

*Q_ST_* was calculated between pairs of the three subspecies under low and high N levels separately. The wild emmer *vs* emmer comparison revealed the effects of primary domestication, the emmer *vs* durum wheat comparison revealed the effects of secondary domestication, and the wild emmer *vs* durum wheat comparison revealed the cumulative effect of domestication. To this end, the model can be reduced to *Y*_*ijl*_ = *S*_*i*_ + *G*_*j*(*i*)_ + ε_*l*(*ij*)_ at each N level. The *Q_ST_* value was calculated as 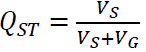 the ratio of between-species and within-species variance.

*Q_ST_* distributions were used to perform a “selection scan” on a restricted number of genes. First, genes were filtered for *H^2^* ≥ 0.7 and, in order not to lose genes whose expression was strongly influenced by N availability, also the species × environment (*S×N*) variance component was evaluated (i.e., every species subgroup × N condition), retaining those genes meeting the threshold *S×N* ≥ 0.2 (Supplementary Figure S1). Successively, we obtained six different *Q_ST_* value distributions (*Q_ST_* _WILD EMMER *VS* EMMER_, *Q_ST_* _EMMER *VS* DURUM WHEAT_ and *Q_ST_* _WILD EMMER *VS* DURUM WHEAT_, each for high and low N conditions) and we retained the 5% upper tail of every distribution. Finally, we compared *F_ST_* and *Q_ST_* values for every gene, discarding *F_ST_* values < 0.01. We confirmed that every retained gene satisfied the condition *Q_ST_* > *F_ST_* allowing it to be classed as undergoing directional selection.

### Differential expression analysis

Differential gene expression was assessed by analysing the pre-processed raw count dataset (32,358 genes). We identified DEGs by comparing (*i*) two conditions (i.e., high and low N levels) within each subspecies, and (*ii*) pairs of the three subspecies under the same N levels, which considered the genotypes nested in species. For the two scenarios, we used three different approaches to detect DEGs: one linear model-based approach implemented in the R package limma (Smyth, 2005), and two Poisson model-based approaches implemented in the R packages edgeR (Robinson et al., 2010) and DESeq2 (Love et al., 2014). In all approaches, the normalization of raw counts was applied by default in the package before differential analysis. To reduce the number of false positives, the intersection of DEGs resulting from the three approaches was retained (Zhang et al., 2014; Rapaport et al., 2013) and the significance threshold was set to an adjusted p-value < 0.001. The DEGs between high and low N levels in at least one subspecies were used for PCA following the *DESeq2* approach (Love et al., 2014), first using all the DEGs, then repeating the analysis on the DEGs considered to be under selection. At each step, counts were normalized using the *vst* method before the *plotPCA* function was applied to define principal components 1 and 2 for the two N levels separately.

### GO enrichment analysis

Enriched terms in the DEGs and genes under selection were identified using agriGO (v.2.0) (Tian et al., 2017) with *T. aestivum* reference annotations and the following parameters: hypergeometric test, multiple hypothesis test adjustment according to the Hochberg FDR procedure at significance level < 0.05 and minimum number of mapping entries of 3.

## Data availability

The raw sequence reads generated and analysed in this study have been deposited in the Sequence Read Archive (SRA) of the National Center of Biotechnology Information (NCBI) under BioProject number PRJNA1015013.

## Acknowledgements

This work was financially supported by Università Politecnica delle Marche, by the PON a3 PlASS Project. PlASS –Platform for Agrofood Science and Safety and by the Transnational Access capacities of the European Plant Phenotyping Network (EPPN, grant n. 284443) funded by the FP7 Research Infrastructures Program of the European Union. H.T. and Z.N. were funded by the European Union’s Horizon 2020 research and innovation program through grant n. 862201.

## Author contributions

R.P. and R.B. conceived and designed the study. T.G. and F.F. carried out the experiments; J.L.D. carried out the library preparation and sequencing. A.P. performed the RNA-Seq analysis. A.P., H.T. and Z.N. performed the bioinformatics analysis and analysed the data. C.D.Q, A.R.L, M.R. provided technical support for RNA-Seq analysis. A.P., R.B., R.P. wrote the paper. Z.N., U.S., V.D.V., G.F., E.Bi., L.N., E.Be., N.P., P.D.V. reviewed and contributed to the editing of the manuscript. All authors have read and approved the manuscript.

## Supplementary Materials

**Supplementary Figure S1:**
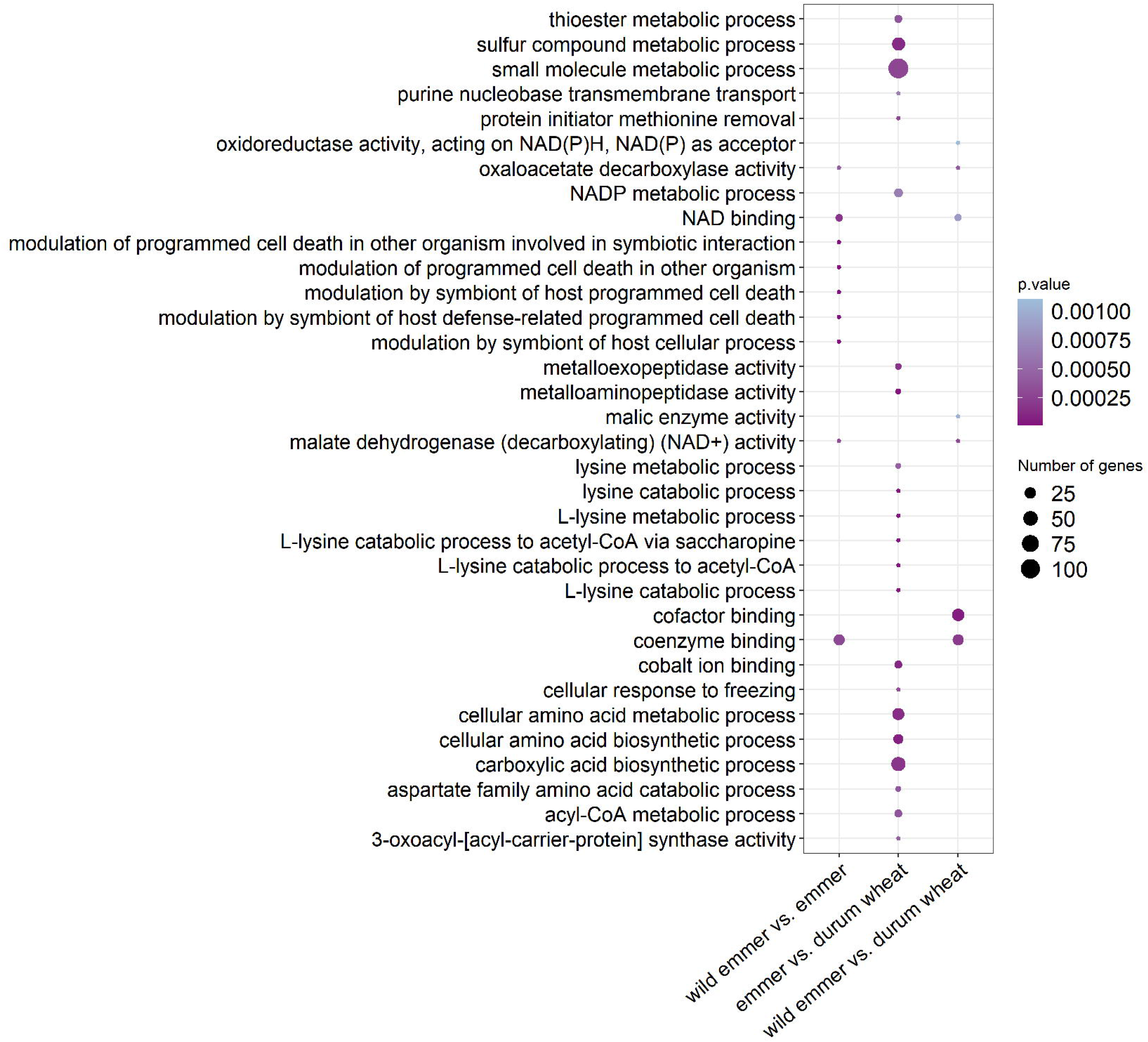
Workflow of gene expression selection scanning.

**Supplementary Figure S2:**
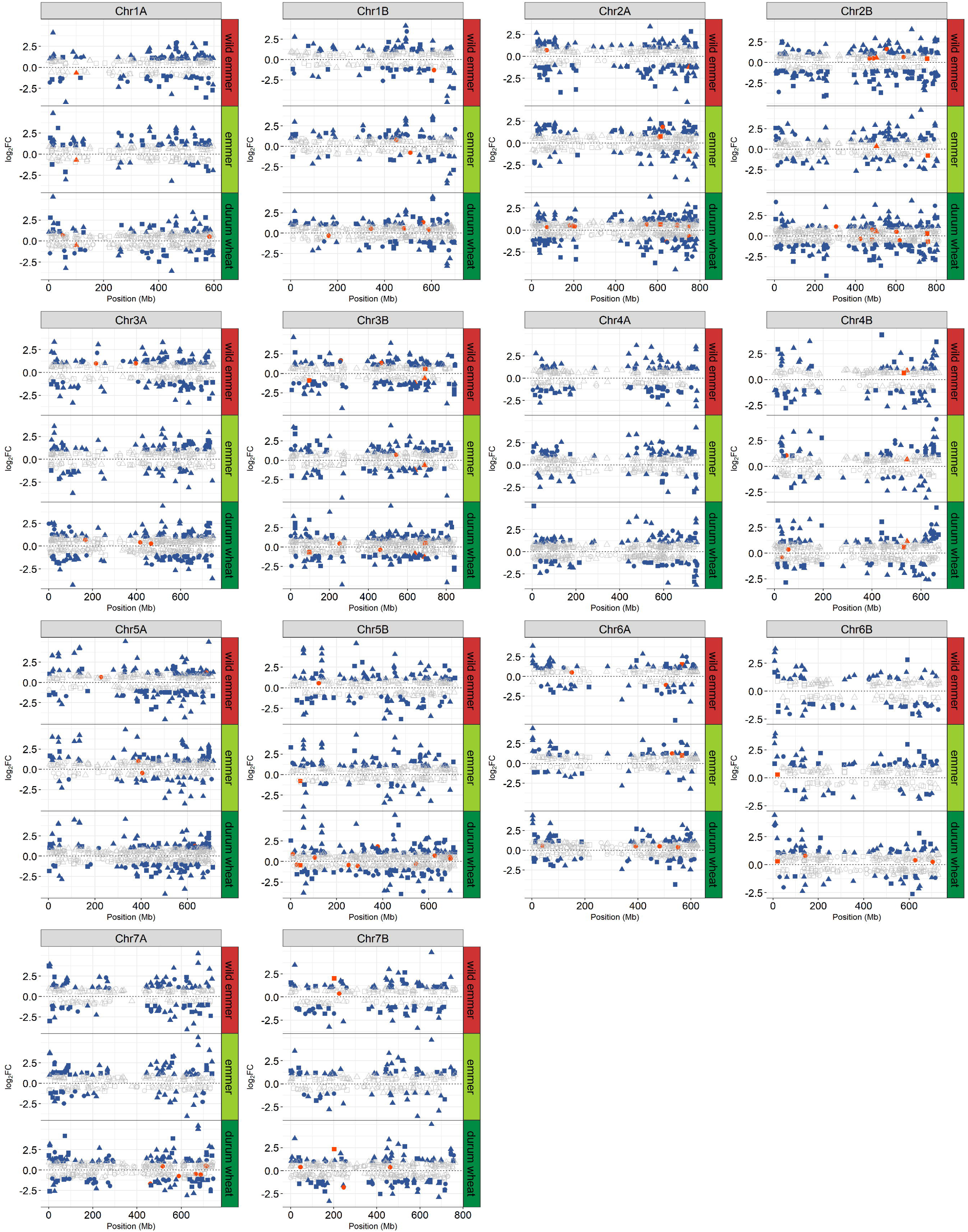
GO categories of genes under selection.

**Supplementary Figure S3:**
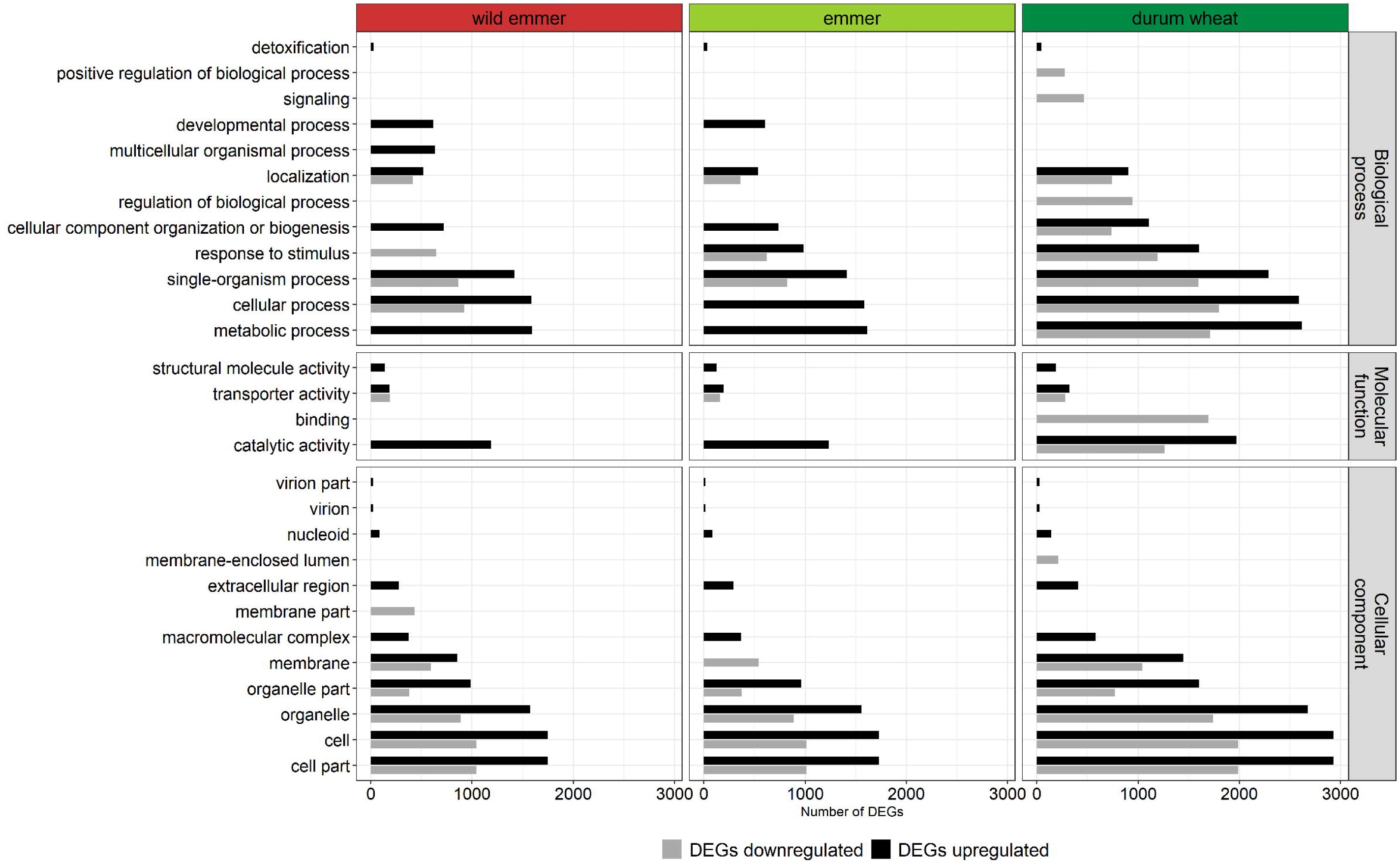
Genome-wide distribution of differentially expressed genes (DEGs) in the comparison between contrasting nitrogen conditions within each subspecies.

**Supplementary Figure S4:**
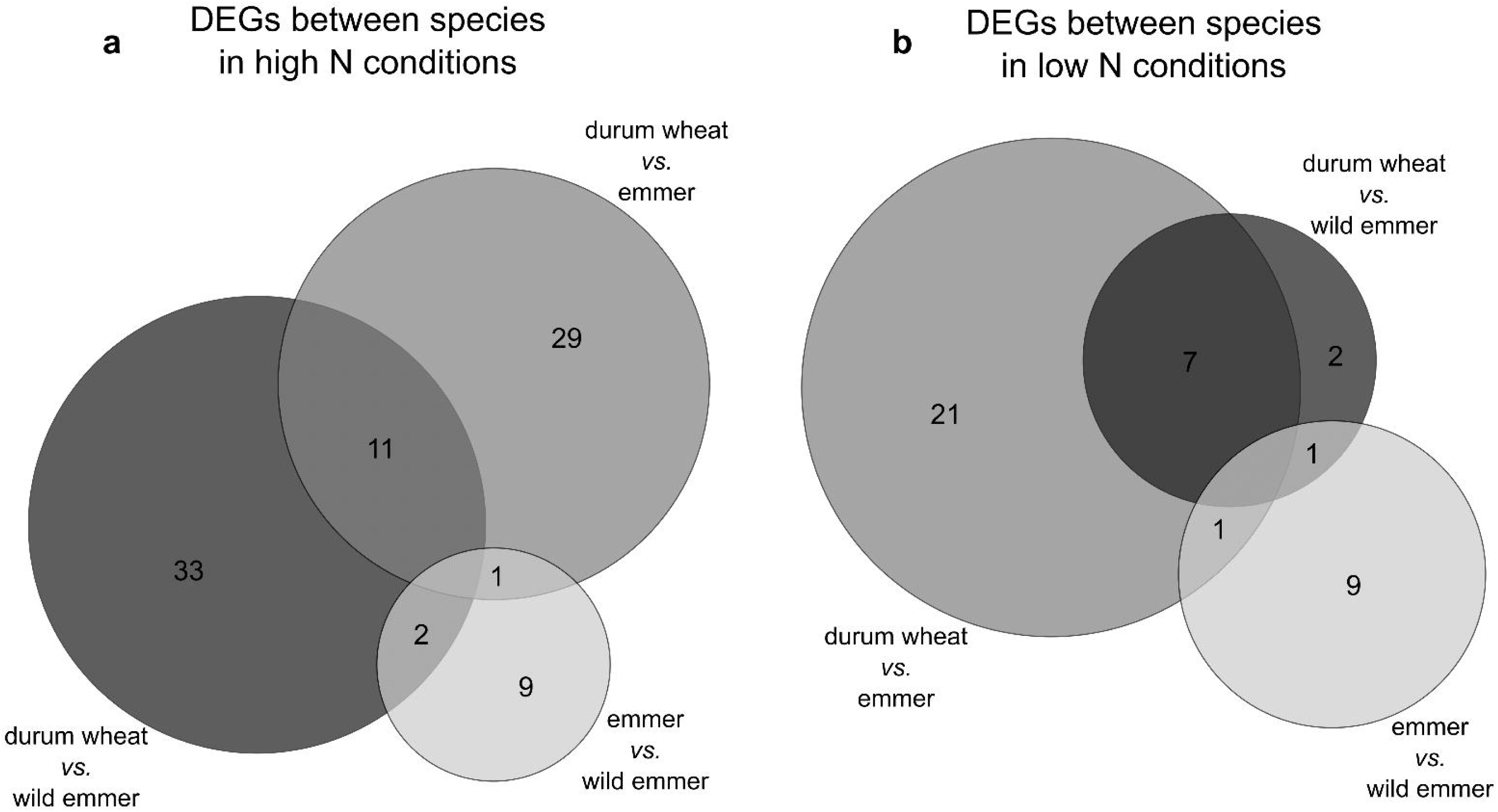
GO classification of differentially expressed genes (DEGs) in the comparison between contrasting nitrogen conditions within each subspecies.

**Supplementary Figure S5:**
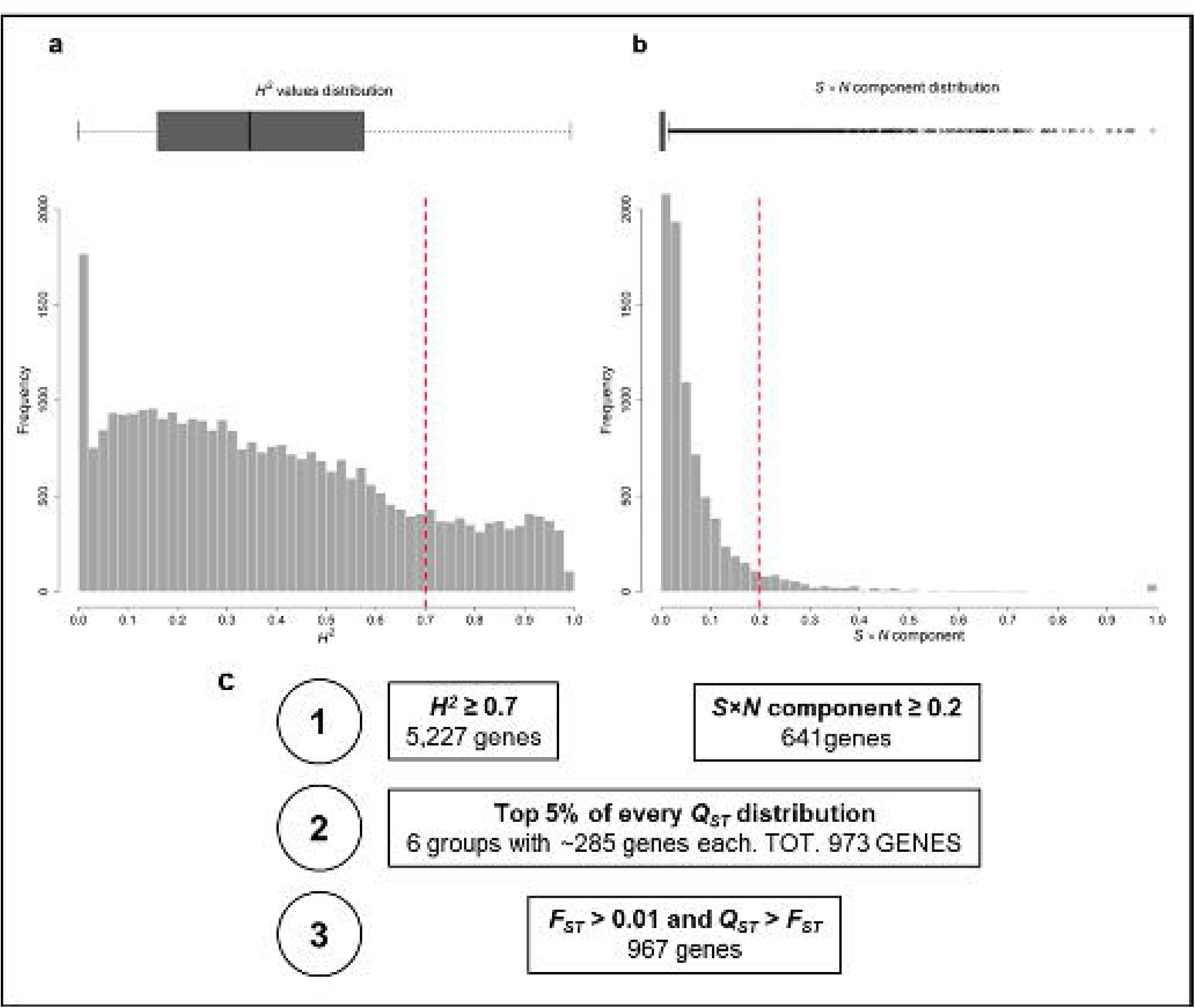
Differentially expressed genes (DEGs) between subspecies.

**Supplementary Table S1:** List of the 128 samples and read mapping results.

**Supplementary Table S2:** Raw read counts of the 32,358 genes.

**Supplementary Table S3:** Mean CV_A_ in gene expression for the three wheat taxa and loss of expression diversity. The loss of expression diversity is shown for two gene subgroups (6,991 DEGs and 25,367 non-DEGs).

**Supplementary Table S4:** List of the 967 genes retained from the “selection scan”. Each gene is accompanied by its functional annotation and the group in which the selection signal was detected.

**Supplementary Table S5:** List of GO “Biological process” and “Molecular function” subcategories for differentially expressed genes (DEGs). GO subcategories are shown for upregulated and downregulated genes under different nitrogen conditions for each subspecies, satisfying the criterion p ≤ 10^−5^.

**Supplementary Table S6:** Functional annotations of the differentially expressed genes (DEGs) between nitrogen conditions in each subspecies. Genes with the top 5% |log_2_FC| values are shown.

**Supplementary Table S7:** Functional annotations of the differentially expressed genes (DEGs) between subspecies under all nitrogen conditions. The corresponding log_2_FC values are shown.

**Supplementary Table S8:** Functional annotation of the 101 genes selected by the integration of selection signatures and differential expression analysis between nitrogen conditions.

